# Platelets are dispensable for the ability of CD8+ T cells to accumulate, patrol, kill and reside in the liver

**DOI:** 10.1101/2021.11.09.467964

**Authors:** James H. O’Connor, Hayley A. McNamara, Yeping Cai, Lucy A. Coupland, Elizabeth E. Gardiner, Christopher R. Parish, Brendan J. McMorran, Vitaly V. Ganusov, Ian A. Cockburn

## Abstract

Effector and memory CD8+ T cells accumulate in large numbers in the liver where they play key roles in the control of liver pathogens including *Plasmodium*. It has also been proposed that liver may act as the main place for elimination of effector CD8+ T cells at the resolution of immune responses. Platelets and the integrin LFA-1 have been proposed to be critical for the accumulation of protective CD8+ T cells in the liver; conversely, asialo-glycoprotein (ASGP) expression on the surface of CD8+ T cells has been proposed to assist in elimination of effector T cells in the liver. Here we investigated the contributions of these interactions in the accumulation of CD8+ T cells activated *in vitro* or *in vivo* by immunization with *Plasmodium* parasites. Using *Mpl*^-/-^ mice with constitutive thrombocytopaenia and antibody-mediated platelet depletion models we found that severe reduction in platelet concentration in circulation did not strongly influence the accumulation and protective function of CD8+ T cells in the liver in these models. Surprisingly, inhibition of ASGP receptors did not inhibit the accumulation of effector cells in the liver, but instead prevented these cells from accumulating in the spleen. We further found that enforced expression of ASGP on effector CD8+ T cells using ST3GalI knockout cells lead to their loss from the spleen. These data suggest that platelets play a marginal role in CD8+ T cell function in the liver. Furthermore, ASGP-expressing effector CD8+ T cells are retained in the liver but are lost from the spleen.

## Introduction

Activated and memory but not naïve CD8+ T cells accumulate in large numbers in the liver [1, 2]. These populations of CD8+ T cells are capable of controlling infections by major liver pathogens including the malaria parasite *Plasmodium* and Hepatitis B virus (HBV) [3-5]. In particular, a population of tissue resident memory cells appears to mediate potent protection against disease [5-8]. CXCR6 expressing tissue-resident memory (TRM) cells patrol the hepatic sinusoids using primarily LFA-1-ICAM interactions to find and eliminate pathogens [7, 9]. The presence of these highly protective CD8+ T cells sheds new light on an older body of literature that suggested that the liver was a “graveyard” for CD8+ T cells [10]. In support of this hypothesis liver-primed CD8+ T cells can be cleared by hepatocytes in a process of emperipolesis [11, 12]. It has further been suggested that this process may be mediated in part by interactions between surface asialo-glycoproteins (ASGPs) and their receptors which are highly expressed by hepatocytes [13-15]. As such the factors affecting the migration of CD8+ T cells in the liver are incompletely understood as are those that predispose to retention vs. apoptosis in this organ.

In addition to LFA-1-ICAM1 interactions it has been proposed that retention of CD8+ T cells in the liver is mediated by interactions with platelets [16]. Platelets are well known to encounter microbes and antigens *via* both innate and adaptive immune processes and help to shape subsequent adaptive responses [17, 18]. Platelets are implicated in the accumulation of neutrophils in the inflamed liver [19, 20]. In a mouse model of hepatitis, it was shown that depletion of platelets leads to enhanced disease due to the accumulation of CD8+ T cells which are pathogenic in this situation [21]. Subsequent work has shown that platelets act as landing pads for CD8+ T cells to dock on prior to establishing residence in the hepatic sinusoids [16]. In contrast, the role of platelets in the clearance of the malaria-causing *Plasmodium* parasite from the liver has not been studied. Protective CD8+ T cells can be induced by vaccination with viral vectors or attenuated parasites [22, 23], but blood stage infection can ablate this vaccine-induced protection [24]. Importantly, fulminant blood stage infection induces thrombocytopaenia in both humans and rodent models of malaria [25-27], thus if platelets were required for the formation and maintenance of resident CD8+ T cell populations in the liver, an acute loss of platelets might be expected to ablate protective CD8+ T cell responses.

Finally, an older body of literature suggests that interactions between ASGPs and their receptors which are abundantly expressed in the liver might mediate accumulation of activated CD8+ T cells in this organ [13, 14]. CD8+ T cells express high levels of ASGPs upon activation [28, 29], which is associated with apoptosis of activated lymphocytes and may be a mechanism of activated lymphocyte removal from the liver, providing a potential explanation for the “graveyard hypothesis” [15, 30]. However, these earlier experiments were performed with neuraminidase treated lymphocytes (a mix of cell types) rather than pure populations of activated CD8+ T cells, so the specific role of ASGP receptors, and the liver more generally, in CD8+ T cell retention and clearance in the liver has not been tested.

To understand how protective memory responses arise in the liver we investigated the roles of platelets and ASGPs in the retention and removal, respectively, of CD8+ T cells from the liver. Counter to findings in the HBV model [16], we found little role for platelets in the accumulation and effector function of CD8+ T cells in the liver in our malaria model. We further found that though the formation of liver-resident memory was associated with the downregulation of ASGPs, interactions with ASGP receptors were not critical for accumulation of effector cells in the liver. These data suggest that LFA-1 is the critical factor for CD8+ T cell accumulation in the liver, and that the spleen rather than the liver may be the main site of effector T cell apoptosis.

## Materials and Methods

### Mice

C57BL/6.J mice, B6 CD45.1, OT-I mice [31], ITGAL-C77F (*Itgal*^-/-^) [7], uGFP [32] and Granzyme B cre mice [33] were bred in-house at the Australian National University (ANU). *Mpl*^-/-^ [34] and *St3gal1*^f/f [35]^ mice were purchased from the Jackson Laboratory. Mice were maintained house under specific pathogen-free conditions except during infection experiments. Mice were aged matched between 6-8 weeks, and were sex matched for all experiments. All animal procedures were approved by the Animal Experimentation Ethics Committee of the Australian National University (Protocol numbers: A2016/17; 2019/36). All research involving animals was conducted in accordance with the National Health and Medical Research Council’s Australian Code for the Care and Use of Animals for Scientific Purposes and the Australian Capital Territory Animal Welfare Act 1992.

### Immunisations, in-vivo platelet depletion and lectin blockade

Mice were immunised intravenously (i.v) with 5×10^4^ *P. berghei* CS^5M^ [36] sporozoites dissected by hand from the salivary glands of *Anopheles stephensi* mosquitos generated in-house within a quarantine approved facility. Prior to immunisation, sporozoites were irradiated at 200kRad of gamma radiation and delivered to each subject. Mice were monitored for the following 21 days for any sign of breakthrough infection both through behavioural changes and blood smear analysis. Platelet depletion of experimental mice was achieved using monoclonal antibodies at a concentration of 20μg per mouse in PBS and delivered i.v via the tail vein: anti-GPIbα (R300 polyclonal-Emfret), anti-αIIbβ3 (Leo.H4-Emfret). Platelet depletion occurred within 30 mins post-injection, however mice were monitored for 60 mins for adverse reactions or excessive bleeding. Control mice received 20μg of an isotype control antibody diluted in PBS and delivered i.v via the tail vein. For lectin blockade, mice were treated with Asialofetuin or Fetuin (Sigma-Aldrich) i.v at concentrations ranging from 1mg/mouse to 3mg/mouse diluted in cold PBS and delivered prior to adoptive transfer studies.

### In vitro activation of T-lymphocytes

Single cell suspensions of C57BL/6 splenocytes were obtained from euthanised animals and incubated with 1μg/ml of SIINFEKL ovalbumin peptide to stimulate T cells. The cells were then co-cultured with a single cell suspension of OT-1 splenocytes in T75 tissue culture flask (ThermoFisher) for 2 days. On day 3, cells were subpassaged into fresh complete RPMI supplemented with 12.5U/ml of rhIL-2 (Peprotech) and incubated for a further 24 hours. The cells were subpassaged a final time with fresh media and IL-2 before being purified on a Histopaque® gradient and transferred.

### Adoptive transfer of T-lymphocytes

OT-I cells were purified on a Histopaque gradient post activation *in-vitro* or eluted from a CD8-negative selection MACS column (Miltenyi) from single cell suspensions of splenocytes. Once purified, the cells were stained (CellTrace™ Violet/ CellTrace™ CFSE) diluted and transferred i.v into sex matched C57BL/6 recipients unless otherwise indicated. For radiation-attenuated sporozoite (RAS) immunisation strategies, 2×10^4^ naïve cells were transferred 24 hours prior to RAS delivery. For intravital imaging and lymphocyte tracking experiments, 5×10^6^ cells were transferred to each mouse. For naïve and activated co-transfer experiments, approximately 2.5×10^6^ cells of each type were transferred. For protection assays, 2×10^6^ cells were transferred 4 hours prior to infection with *P*.*berghei*.

### Lymphocyte harvesting and flow cytometry

Single cell suspensions were isolated from euthanised mice and prepared using specified protocols to isolate cells from the liver, lung, spleen, lymph node and bone marrow. Single cell suspensions were incubated using Fc-Block (Biolegend) for 15 minutes on ice followed by staining with fluorescently conjugated Abs: anti-CD11a (clone M17/4- Biolegend), anti-KLRG1 (clone MAFA- Biolegend), anti-CD69 (clone H1.2F3- Biolegend), anti-CD8 (clone 5H10-1/clone 53.67- Biolegend), anti-Ly5a (clone A20- Biolegend), anti-Ly5b (clone 104- Biolegend), anti-CD62L (clone MEL14- Biolegend), anti-Vα2 (clone B20.1- Biolegend), anti-CD3 (clone 17.A2- Biolegend). Samples were then resuspended in FACs buffer with viability dye (7-AAD) and transferred to cluster tubes for cytometric analysis. If cells were to be fixed, processing omitted the 7-AAD step and incubated with a fixable live dead dye prior to incubation with fixation buffer (Biolegend) (15 mins). Cells were analyzed using a LSRII flow cytometer (Becton Dickinson) or Fortessa X20 cytometer (BD Biosciences). Data were analysed using FlowJo analysis software (Tree Star).

### Platelet isolation and lymphocyte co-culture

Blood was collected *via* tail vein bleed and collected in acid citrate dextrose solution (ACD). The blood was then stored at room temperature and centrifuged (250 g, 16 mins, 21°C). The upper platelet-rich plasma (PRP) layer was removed and transferred to a new tube and rested for 15 minutes at room temperature. The PRP was then centrifuged (1200 g, 5 mins, 21°C) and the pellet was resuspended in platelet wash buffer (150mM NaCl containing 10mM trisodium citrate and 1% (w/v) dextrose, pH 7.4) and rested for 15 minutes. The suspension was centrifuged again (1200 g, 5 mins, 21°C) and gently resuspended in Tyrode’s buffer to a concentration of 2×10^8^ platelets/ml before being rested at room temperature for 60 mins then 50μl transferred to wells containing 1×10^6^ OT-I lymphocytes. Wells were cultured for 1 hour in a 10:1 ratio prior to being washed and stained with fluorochrome-conjugated Abs: anti-CD41 (clone MWReg- Biolegend) and anti-CD8 (clone 53.67- Biolegend). Cultures were then analysed using Amnis ImageStream®X (Merck Millipore).

### Assessment of parasite burden

Parasite burden was measured via qRT-PCR using primers that recognise *P*.*berghei* specific sequences within the 18S rRNA and SYBR Green (Applied Biosystems) as outlined previously [37]. Parasite burdens were normalised with GAPDH expression.

### Multiphoton microscopy

Mice were prepared for microscopy *in vivo* as described previously (van de Ven *et al* 2013). Once the mouse was ready and applied to the movable platform of a Fluoview FVMPE-R multiphoton microscope, the platform was raised to ensure contact of the XLPLN25XWMP2 objective lens with a drop of water on the coverslip (25x, NA1.05, water immersion; 2mm working distance). For the analysis of motility of cells activated *in vitro*, a 50μm Z-stack (2μm/slice) was typically acquired using the galvo-scanner at a frame rate of typically 2 frames per minute. For naïve and activated motility analysis, a single slice was acquired using the resonant scanner with 3-6x averaging at a rate of approximately 3 frames/second. The images were acquired using the FV30 software (Olympus) and exported to Imaris (Bitplane) for track analysis using autoregressive motion algorithm and polarity analysis.

### Statistical analysis

Data is shown as individual data points with bars (where shown) indicating mean ± S.D.. Data from two or more experiments were analysed using linear mixed modelling (LMM) in R libraries *lm4* and *nlme* (The R Foundation for Statistical Computing). In the instance of data being pooled from several experiments, each experiment was included as a random effect blocking factor in the LMM analysis. Cellular data was log transformed where data is presented on a log scale, prior to statistical analysis. For all other cellular data where experimental blocking factors did not need to be accounted for, analysis was conducted in GraphPad Prism v7.

## Results

### Activated but not naïve CD8+ T cells accumulate and patrol in the liver sinusoids

To determine the different homing and migration patterns of activated and naïve CD8+ T cells we co-transferred differentially labelled *in vitro* activated and naïve OT-I T cells specific for the SIINFEKL epitope from chicken ovalbumin to naïve mice (Figure 1A). Preliminary studies showed that, similar to *in vivo* activated cells, *in vitro* activated cells had elevated levels of LFA-1 and could be labelled with Peanut Agglutinin (PNA) which binds ASGPs and to a lesser extent asialo-gangliosides such as asialo ganglio-N-tetraosylceramide (asialo GM) [38] (Figure S1 A and B). We also determined that effector cells were able to bind platelets at a ∼10-fold higher frequency than naïve cells as revealed by CD41/CD8 co-staining (Figure S1C). Finally, Imageflow analysis revealed that activated cells typically bound multiple platelets while the few naïve cells that bound platelets only bound a single platelet (Figure S1D).

**Figure 1.**
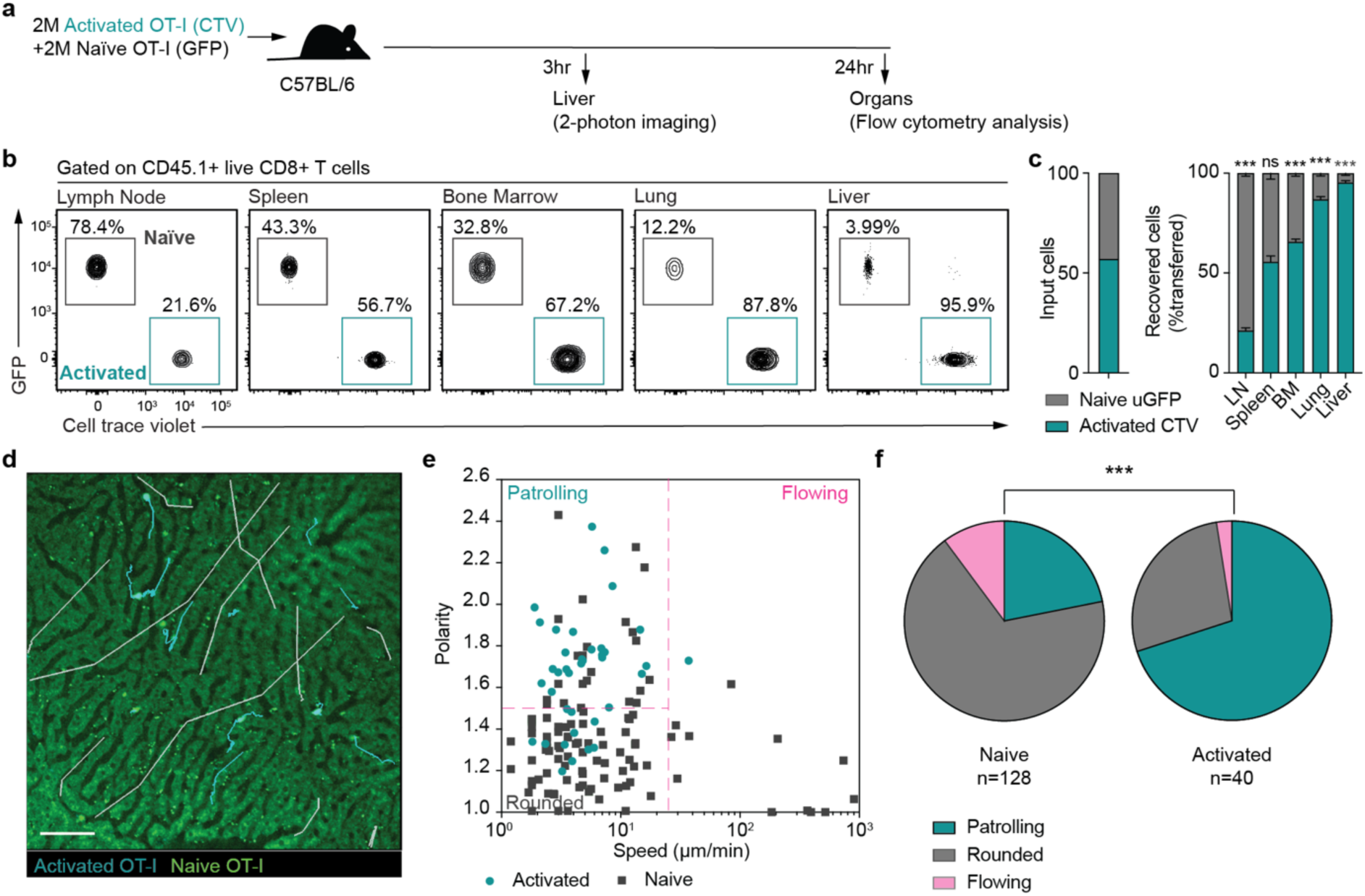
*In vitro* activated CD8 T-lymphocytes demonstrate enhanced migration to organs such as the liver and develop an effector phenotype with patrolling behaviour. (A) 2×10^6^ SIINFEKL pulsed OT-I T-cells (CTV) and 2×10^6^ naïve GFP^+^ OT-I cells were transferred to C57BL/6 mice. 4 hours post adoptive transfer, flow cytometry analysis was conducted on axillary lymph nodes, spleen, bone marrow, lung and liver from recipients. (B-C) Proportion of donor cells isolated from each organ after co-transfer of equal amounts of activated (blue) and naïve (green) OT-I cells; data in B-C from 5 mice per group in one of two independent experiments analyzed via one-sample t test; bars are mean ± S.D; *** p<0.001 (D) Livers of recipient mice upon 2-photon microscopy using resonance scanning to collect time-lapse movement at 3 frames per second demonstrating elongated linear tracks of naïve cells (white) compared to short repetitive tracks of activated patrolling cells (blue). (E) Mean speed versus polarity of activated (blue) and naïve (black) T-lymphocytes in the liver. (F) Proportion of both naïve and *in vitro* activated cells exhibiting different T-cell migration behaviours in recipient mice post transfer. Scale bar is 50μm. Data in D-F is pooled from 2 independent experiments analyzed by χ^2^ test.

Twenty-four hours after the co-transfer of GFP+ naïve and cell trace violet (CTV)-labelled activated OT-I cells, total lymphocytes were recovered from the liver, lung, bone marrow, spleen and lymph nodes and analysed by flow cytometry. Naïve and activated cells accumulated roughly evenly in the spleen while naïve cells specifically accumulated in the lymph node (Figure 1B); conversely activated CD8+ T cells preferentially accumulated in the liver and lung, and to a lesser extent in the bone marrow (Figure 1C). We further examined the behaviour of the co-transferred naïve cells and activated cells within in the liver by multi-photon microscopy. In these studies, we used a resonance scanner to take high frame rate movies enabling us to capture both crawling cells and faster flowing cells in the blood stream (Movie S1; Figure 1D). In agreement with our previous analysis [7], activated CD8+ T cells undertook a crawling behaviour in the liver sinusoids in which they become elongated and move both with and against the blood flow at average speed of <25μm/min (Figure 1D-E), while naïve cells were generally observed to be either flowing in the blood or rounded up and stationary (Figure 1E-F), a phenotype which we have previously associated with activated *Itgal*^*-/-*^ OT-I cells that lack expression of the LFA-1 integrin [7].

### Platelets are not required for CD8+ T cell effector function in the Plasmodium infected liver

To test the role of platelets in CD8+ T cell effector function in the liver we used *Mpl*^-/-^ mice, which carry a mutation in the thrombopoetin receptor (*Mpl*) gene and have around 15% of the normal number of circulating platelets [34]. We transferred *in vitro* activated OT-I cells to wild-type and platelet deficient *Mpl*^-/-^ mice that were subsequently infected with *P. berghei* CS^5M^ parasites. *P. berghei* CS^5M^ parasite express the SIINFEKL epitope recognized by OT-I cells within the surface circumsporozoite protein [36]. Parasite burden was subsequently measured by RT-PCR [37]. In both WT and *Mpl*^-/-^ mice, activated OT-I T cells conferred significant protection against infection, however the degree of protection was significantly lower in *Mpl*^-/-^ mice suggesting that platelets may play a role in protection (Figure 2A).

**Figure 2.**
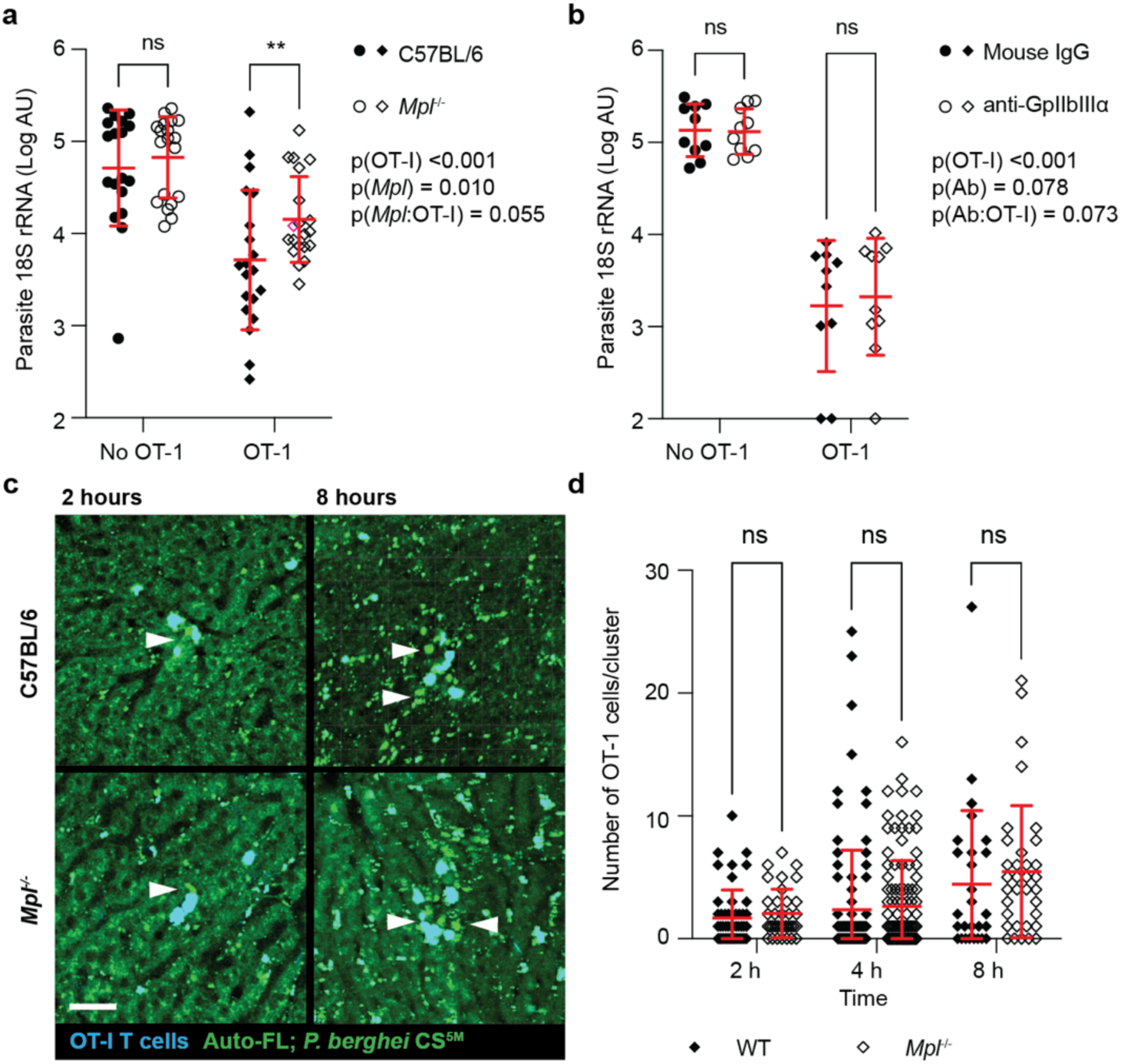
OT-I effector lymphocyte killing capacity of *Plasmodium* is minimally affected by platelets *in vivo*. WT and *Mpl*^*-/-*^ recipients (A) and platelet-depleted recipients (B) received 5×10^6^ *in vivo* activated OT-I cells. 24 hours later, recipients were infected with 5×10^3^ *Plasmodium* sporozoites. *Plasmodium 18S* rRNA levels were measured 24 hours post infection to assess protective function of adoptively transferred OT-I cells; data in A is pooled from 4 similar experiments and data in B is pooled from 2 similar experiments with 4-5 mice per group analysed via LMM; bars are mean ± S.D.; ** p<0.01. (C) WT and *Mpl*^*-/-*^ recipient mice received 5×10^6^ OT-I cells (blue) and were infected with GFP expressing *P. berghei* CS^5M^ (green-white arrow); scale bar 50 μm. (D) Using intravital imaging, the number of cells surrounding each parasite within the liver was assessed at 2, 4 and 8 hours post infection in WT and *Mpl*^*-/-*^ recipients; data from at least 3 mice per group per timepoint; bars are mean ± S.D.; analysed via LMM.

However *Mpl*^-/-^ mice have elevated levels of circulating thrombopoietin which may alter haematopoiesis in these mice potentially affecting the protective capacity indirectly [39]. To specifically investigate the role of platelets, we also measured the ability of activated CD8+ T cells to protect mice that had undergone platelet depletion using an anti-αIIbβ3 mAb, which targets integrin αIIbβ3 expressed only on the platelet membrane, and, in our hand, results in >97% reduction in the numbers of circulating platelets 2 hour post injection [40]. In this system CD8+ T cells transferred to platelet-depleted mice were able to protect against malaria challenge equally well as cells transferred to platelet replete/sufficient/intact mice (Figure 2B).

CD8+ T cell killing is preceded by the formation of clusters of activated CD8+ T cells around the infected hepatocyte [41-43]. We therefore examined the kinetics of cluster formation 2, 4 and 8 hours after transfer of CD8+ T cells to infected *Mpl*^-/-^ and WT mice. However, no difference in the size of clusters formed around infected hepatocytes in *Mpl*^-/-^ mice compared to wild-type controls was detectable by quantitative microscopy suggesting that platelets are not required for the localization of *Plasmodium*-infected hepatocytes by CD8+ T cells (Figure 2C-D). Overall, these data suggest that platelets play limited roles in CD8+ T cell accumulation within the liver and in the control of infection once cells are established in the liver.

### Platelets are not required for normal CD8+ T cell motility and accumulation in the liver

Earlier studies describing a role for platelets in the homing of CD8+ T cells in the liver suggested that platelets were required for the initial tethering of activated CD8+ T cells to the walls of the hepatic sinusoids [16]. We hypothesised, therefore, that platelets may be acting earlier than the timepoints examined in the above experiments. Activated CD8+ T cells were therefore transferred to *Mpl*^-/-^ and control mice and CD8+ T cell accumulation was measured in the liver and spleen (Figure 3A). We further considered that the tethering effect of platelets might be partially redundant with LFA-1 binding by CD8+ T cells in the liver. We therefore also activated *Itgal*^-/-^ OT-I cells and co-transferred these with activated wild-type OT-I cells. Surprisingly, after 20 mins *Itgal*^-/-^ cells were observed to accumulate in greater numbers within the liver than wild-type cells (Figure 3B-C), however, they were then rapidly lost from this organ and subsequently accumulated in the spleen (Figure 3D-E). Despite clear differences in the accumulation of *Itgal*^-/-^ versus wildtype cells, the kinetics of accumulation of both *Itgal*^-/-^ and wild-type OT-I cells as suggested by linear mixed effect modelling was similar between wild-type and *Mpl*^-/-^ hosts regardless of which organ was studied (Figure 3B-E). Overall, we were unable to discern any defect in CD8+ T cell accumulation in the livers of *Mpl*^-/-^ mice, although we were able to confirm a role for LFA-1 in this process.

**Figure 3.**
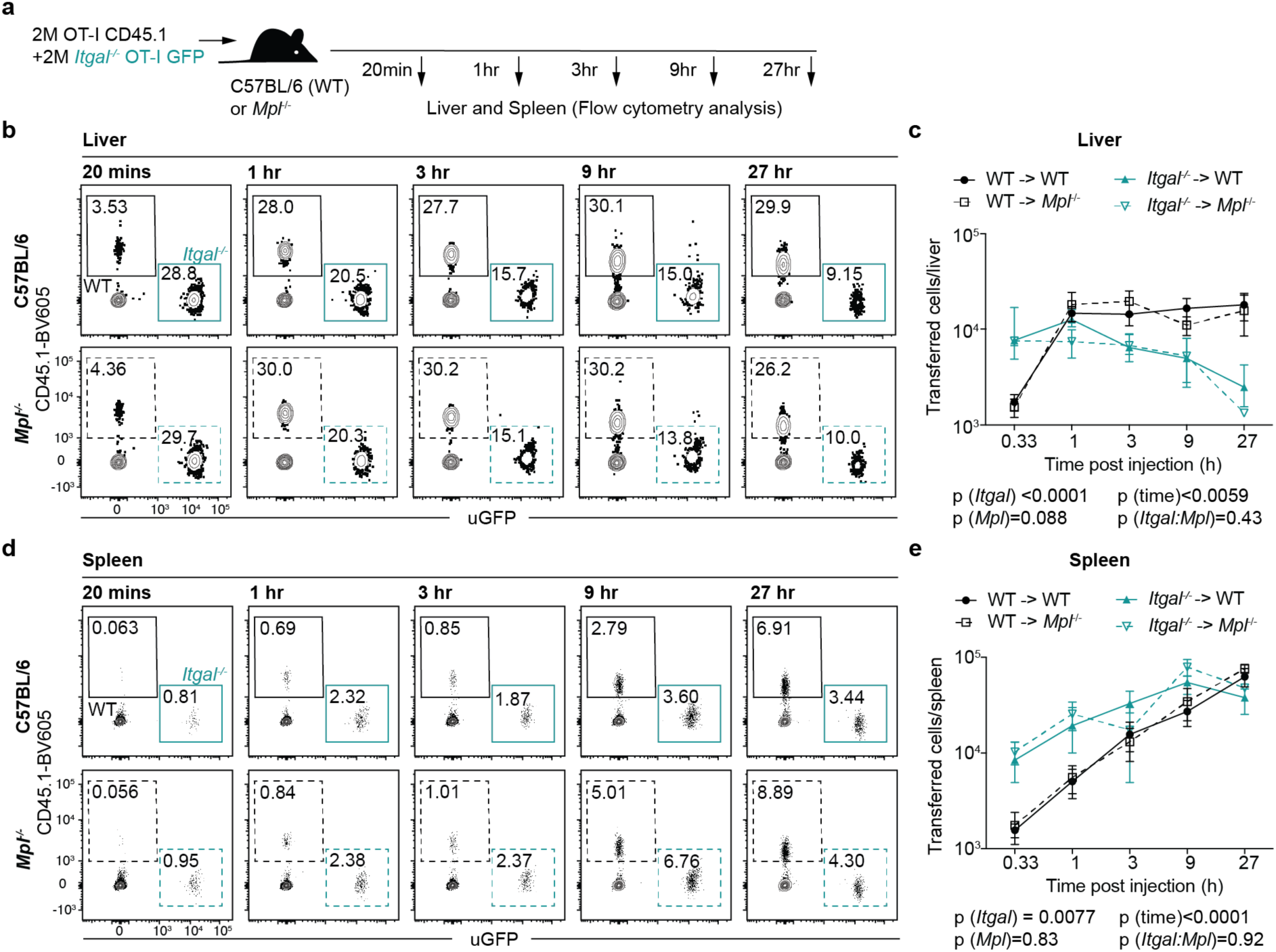
LFA-1 binding acts as a dominant homing mechanism of OT-1 cells from the circulation to the spleen and liver. (A) 2×10^6^ OT-I cells, and 2×10^6^ OT-I GFP^+^ *Itgal*^*-/-*^ cells were co-transferred to C57BL/6 or *Mpl*^*-/-*^ recipients. Single cell suspensions from the liver and spleen were prepared at several timepoints post transfer (20m, 1hr, 3hr, 9hr and 27hr) and flow cytometry was used to assess *Itgal*^-/-^ (blue) and WT (black) donor cell accumulation in the liver (B-C) and spleen (D-E) of C57BL/6 (solid line) and *Mpl*^*-/-*^ (dashed line) recipients. Results are pooled from two independent experiments with 5 mice per group; bars are mean ± S.D; analyzed via LMM.

To determine if there were any differences in the behaviour of cells once retained in the liver, activated CD8+ T cells were transferred to wild-type or *Mpl*^-/-^ mice and the patterns of migration in the sinusoids 4 or 24 hours after transfer was measured using multiphoton microscopy. Using this analysis, we were unable to discern any difference in speed, straightness or time spent moving between CD8+ cells in the livers of *Mpl*^*-/-*^ or control animals (Movie S2; Figure S2A-B).

As *Mpl*^-/-^ mice have sufficient residual platelets (∼15% of the normal number) to confer nearly normal haemostatic function, we examined the accumulation of wild-type and *Itgal*^*-/-*^ deficient CD8+ T cells in anti-GPIbα antibody treated mice, which depletes platelets to <5% of normal levels (Figure 4A-B). This same antibody was used in previous studies examining hepatitis B virus-specific CD8+ T cells [16]. Importantly, anti-GPIbα antibodies deplete platelets by inducing the expression of neuraminidase on platelets leading to the exposure of asialo-glycoproteins on the surface of platelets and their clearance by ASGP receptors on hepatocytes [44]. Similar to our results with *Mpl*^-/-^ animals, activated wild-type OT-I T cells accumulated at similar levels in the livers of anti-GPIbα antibody-treated mice and control animals (Figure 4C-D). However, the accumulation of *Itgal*^-/-^ cells in the liver, which was already limited, was further impaired by platelet depletion. Thus, under these conditions we were able to identify a modest role for platelets in the accumulation of activated *Itgal*^-/-^ cells, although the effect of LFA-1 deficiency was much greater (Figure S3B and C). Interestingly, both wild-type and *Itgal*^-/-^ OT-I cells were inhibited in their ability to accumulate within the spleen in the anti-GPIbα antibody treated mice (Figure 4C-D). This contrasted with the lack of phenotype observed in the platelet-deficient *Mpl*^-/-^ mice. This difference could be due to an insufficient loss of platelets in the *Mpl*^-/-^mice, or an off-target effect of the depleting antibodies.

**Figure 4.**
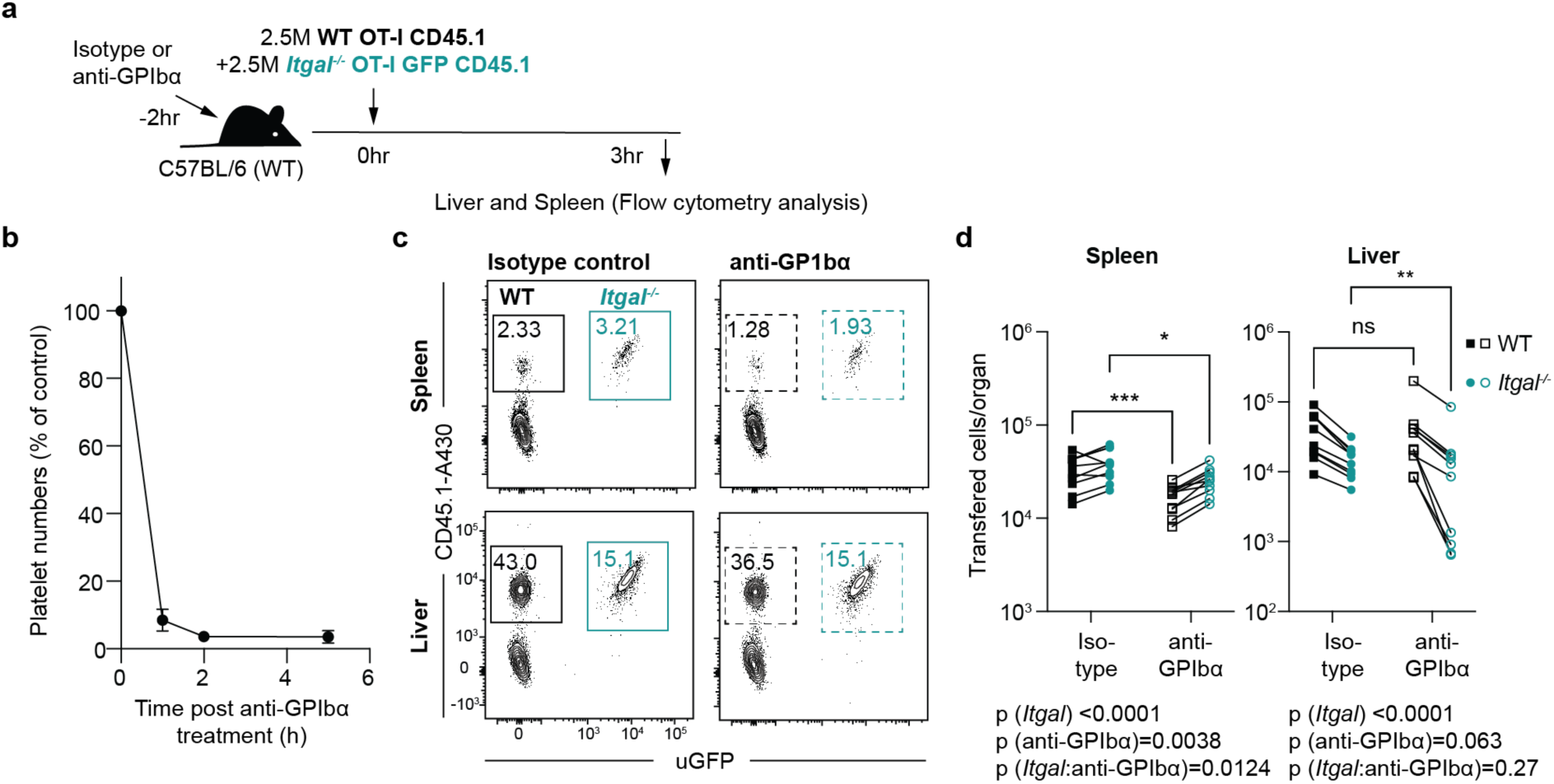
Platelet depletion decreases lymphocyte homing to the spleen however has marginal effect on homing to the liver in the absence of LFA-1. (A) 2.5×10^6^ OT-I cells, and 2.5×10^6^ OT-I GFP^+^ *Itgal*^*-/-*^ cells were co-transferred to C57BL/6 recipients which had received platelet depletion antibody treatment (anti-GPIbα) or isotype control. Single cell suspensions from the liver and spleen of the recipients were prepared at 3 hours post adoptive transfer for analysis. (B) Kinetics of platelet depletion following treatment anti-GPIbα. (C) Representative flow cytometry plots and summary data (D) of the numbers of *Itgal*^-/-^ (blue) and WT (black) donor cell cells in the spleen and liver C57BL/6 (solid line) and anti-GPIBα (dashed line) recipients. Results are pooled from two independent experiments (n=5 per experiment) analyzed via LMM, * p<0.05, ** p<0.01.

### Platelet deficiency does not affect the formation of liver resident memory cell populations

The previous experiments were performed with *in vitro* activated cells which may not fully replicate all aspects of infection or immunization situations. We therefore asked whether platelets may play a role in the formation of T cell populations in the spleen and liver after *in vivo* immunization. In particular, since we have previously identified roles for LFA-1 and CXCR6 in the formation of liver tissue resident memory T cells (TRM) we wished to examine this population [7, 9]. Accordingly, we assessed the ability of *P. berghei* CS^5M^ sporozoite primed mice to form CD8+ TRM in the livers of *Mpl*^-/-^ mice or platelet-depleted mice that had received OT-I cells (Figure 5A; Figure S3A). However, CD8+ T cell populations appeared normal in terms of numbers and phenotypes in *Mpl*^-/-^ mice compared to wild-type mice suggesting that low platelets do not affect the accumulation or maintenance of TRM populations in the liver (Figure 5B-C). In further support of this, administration of the anti-GPIbα antibody 24hr before tissue harvesting also had no effect on the numbers of CD8+ TRM cells in the livers of sporozoite immunized mice (Figure S3B-C).

**Figure 5.**
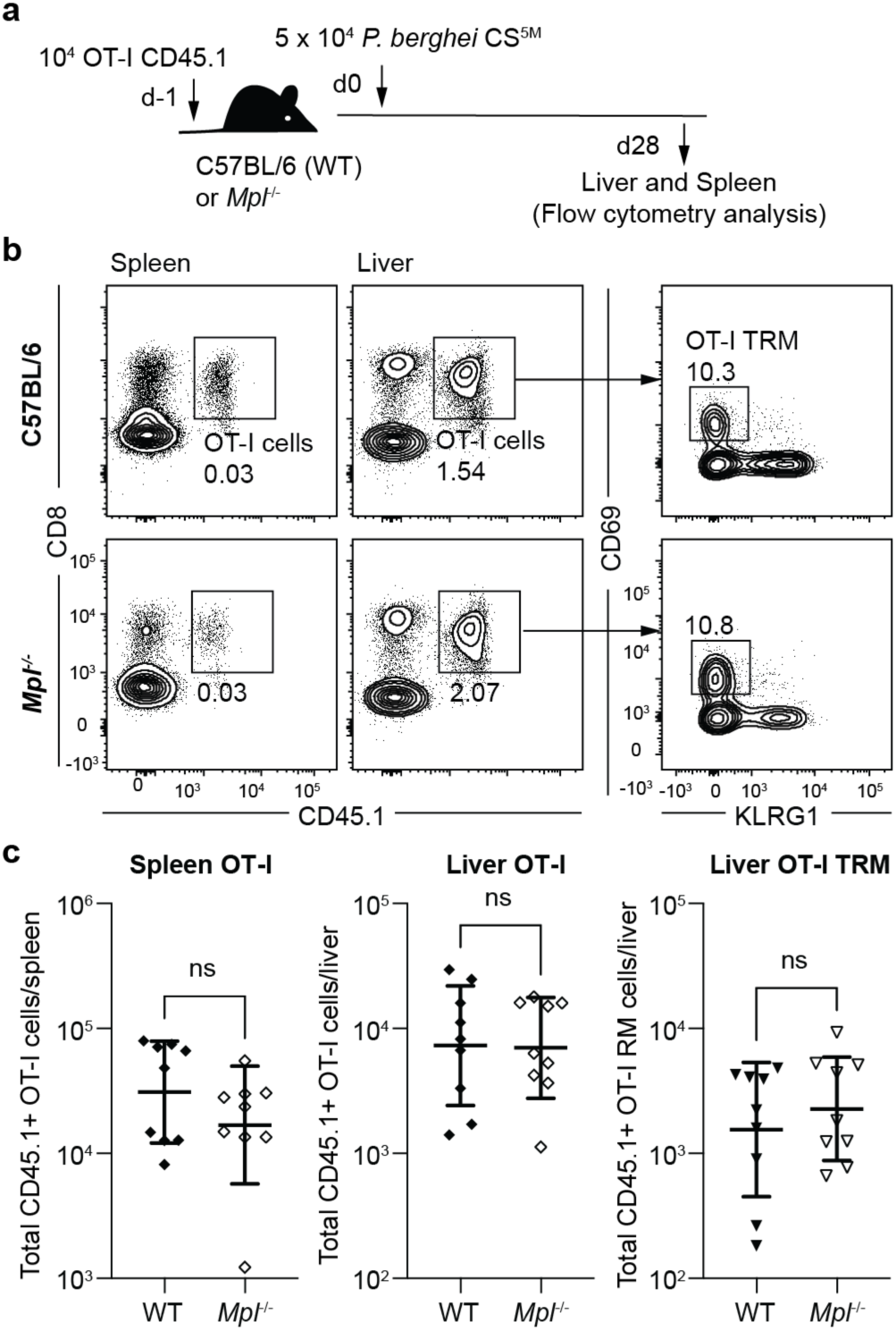
*In vivo* generation of tissue resident memory T cells is not affected by the absence of platelets in the spleen and liver. (A) C57BL/6 and *Mpl*^*-/-*^ recipients received 1×10^4^ OT-I cells prior to immunisation with 5×10^4^ *P*.*berghei* CS^5M^ RAS 24h later to generate CD8 tissue resident memory (TRM) populations *in vivo* after 28 days. Total transferred cells recovered from the spleen and liver were assessed at day 28 post immunisation from both C57BL/6 and *Mpl*^*-/-*^ recipients. (B) Representative flow cytometry plots and (C) summary data pooled from 2 independent experiments each with 5 mice per group; bars are mean ± S.D; analyzed via LMM.

### Asialylated glycoproteins (ASGPs) mediate effector CD8+ T cell accumulation in the red pulp of the spleen

In addition to platelets, ASGPs have been proposed to mediate the accumulation of lymphocytes in the liver. This accumulation is hypothesised to precede the destruction of cells either via apoptosis or via the uptake of cells by hepatocytes also known as emperiopolesis [11, 12, 15, 45]. To investigate the role of ASGPs in the accumulation of activated CD8+ T cells in the liver, *in vitro* activated effector cells were transferred to mice that had received asialo-fetuin (ASF) to block ASGP receptors. Control mice received fetuin which is abundantly glycosylated with sialylated carbohydrates (Figure 6A). Strikingly, ASF did not alter effector T cell accumulation in the liver, but similar to anti-GPIbα treatment, blocked accumulation in the spleen (Figure 6B-C). Because anti-GPIbα treatment results in the release of neuraminidase which desialylates glycoproteins we speculated that this treatment may also be impacting upon effector T cell accumulation in the spleen via blockage of ASGPs. We therefore repeated the ASF blockade experiment including additional groups treated with anti-GPIbα antibodies. However, platelet depletion did not further enhance the exclusion of cells from the spleen via ASF (Figure 6B-C). These data suggest that anti-GPIbα antibody treatment was not inhibiting CD8+ T cell accumulation in the spleen as a result of platelet depletion but rather by ablating of ASGP receptor (ASGPR) function.

**Figure 6.**
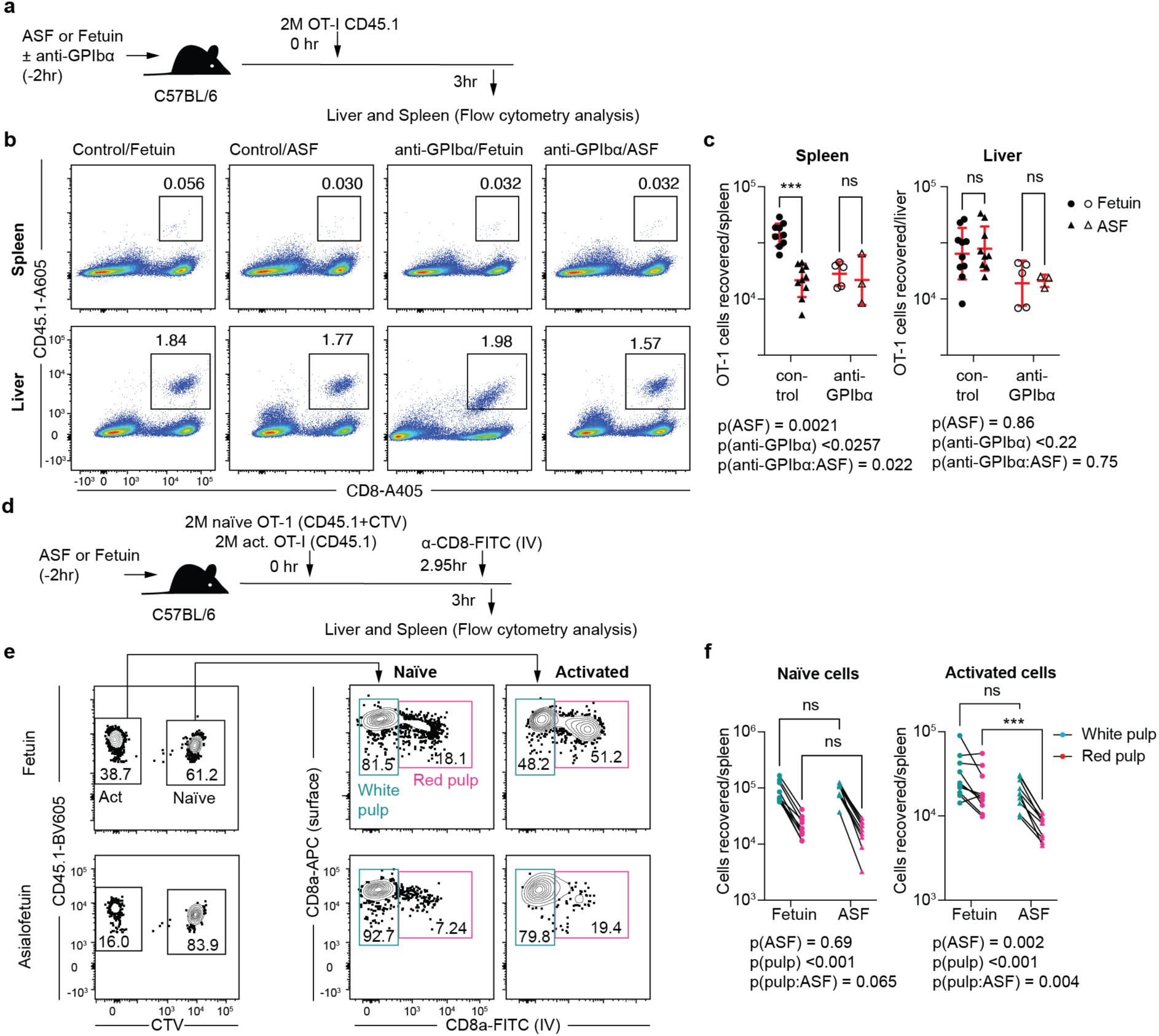
Asialylated glycoprotein residues reduce effector lymphocyte homing to the heavily vascularised red pulp of the spleen, with no effect on homing to the liver. (A) Prior to adoptive transfer of 2×10^6^ OT-I cells, recipient C57BL/6 mice were treated with glycoproteins (ASF or the control Fetuin) and either platelet depletion antibody therapy (anti-GPIbα), or an isotype control. (B) Single cell suspensions from the liver and spleen of the recipients were prepared at 3 hours post adoptive transfer and flow cytometry was used to quantify WT (black) donor cell accumulation in the spleen and liver. (C) Total recovered OT-1 cells from the spleen and liver after antiplatelet therapy (anti-GPIbα /isotype) and glycoprotein blocking treatment (ASF/Fetuin); data are pooled from 2 experiments, one with just non-platelet depleted animals and one with all four groups; bars are mean ± S.D; analyzed via LMM; *** P<0.001. (D) Prior to co-transfer of 2×10^6^ *in vivo* activated OT-I cells and 2×10^6^ naive OT-I cells, recipient C57BL/6 mice were treated with glycoprotein blocking treatment (ASF or the control Fetuin). Immediately prior to culling at 3 hours post transfer, recipients received FITC conjugated anti-CD8 antibody i.v to label CD8+ T-cells flowing in the systemic circulation. (E-F) Single cell suspensions from the spleen were obtained and quantified using flow cytometry. Activated and naïve transgenic CD45.1 cells were isolated and identified as being resident in the red pulp (pink) or white pulp (green) based on FITC labelling *in vivo*. Results are pooled from two independent experiments analyzed via LMM; ***p<0.001. Asterisks (*) indicate significant differences between groups (* p <0.05). ns, not significant.

To further dissect the effect of ASF on effector cell homing to the spleen we examined the specific locations in the spleen in which effector T cells accumulated and determined whether ASF treatment specifically affected migration to particular sub-compartments. Accordingly, we designed an experiment in which naïve CD45.1 OT-I cells and activated OT-I cells were transferred to mice in the presence of ASF or fetuin. Three minutes prior to euthanasia the mice were injected i.v with anti-CD8a antibody (Figure 6D) which labels the cells in the red pulp that are exposed to the circulation, but not those in the white pulp that are shielded from the circulation. As expected, the naïve cells preferentially accumulated in the white pulp and this migration was not affected by ASF treatment (Figure 6E-F), which is consistent with the lack of ASGPs on the surface of naïve CD8+ T cells (Figure S1). However, activated cells accumulated equally between the red and white pulp in the control mice, but were specifically excluded from the red pulp in the ASF treated animals (Figure 6E-F). Thus, interactions with ASGPs appear to mediate the accumulation of effector T cells in the red pulp of the spleen.

### Forced expression of ASGPs drives the loss of effector T cells from the spleen but not the liver

To extend the finding *in vitro* that ASGP expression mediated the accumulation of activated effector T cells in the spleen to an *in vivo* immunization model, we transferred OT-I cells to mice which were then immunized with *P. berghei* CS^5M^ sporozoites. The expression of ASGPs on activated CD8+ T cells was measured using PNA staining on days 7, 14 and 28 post immunization. As expected, large numbers of cells accumulated in the spleen and liver (Figure S4A) and populations of effector (Teff; KLRG1^hi^, CD62L^lo^), effector memory (TEM; CD62L^lo^, CD69^-^, KLRG1^lo^) and central memory cells (TCM; CD62L^hi^, KLRG1^lo^) could be identified in both organs, while the liver also carried substantial numbers of CD69^hi^ CD62Llo KLRG1lo liver TRM cells (Figure S4B-C). In contrast PNA binding was highest on Teff cells at early time points declining with time (Figure S4D-E). PNA binding was also highest initially in the liver but declined to similar levels in the spleen and liver by day 28 (Figure S4D-E). Interestingly TRM cells in the liver had significantly lower PNA binding that other populations suggesting that the establishment of this memory population may be dependent on the loss of ASGP receptors (Figure S4D-E).

To investigate the effects of ASGP expression on CD8+ T cell fate we created a situation in which ASGP expression was enforced on responding CD8+ T cells. Sialylation of Core1 glycoproteins is mediated by the enzyme ST3 beta-galactoside alpha-2,3-sialyltransferase 1 (ST3Gal1; Figure 7A). We therefore crossed our OT-I mice to *St3gal1*^f/f^ x Granzyme B Cre (GzbCre) animals such that ST3Gal1 was deficient in all responding CD8+ T cells and PNA-binding was enforced uniformly (Figure 7B). We used a granzyme B Cre as previous studies with a CD4 Cre showed that enforced expression of PNA in the thymus led to apoptosis and the loss of peripheral CD8+ T cells [35]. In these experiments *St3Gal1*^f/-^ x GzbCre (Het) or KO *St3Gal1*^f/f^ x GzbCre (KO) Ly5B OT-I cells were co-transferred to wild-type (Ly5A) mice with Ly5AB OT-I wild-type cells prior to immunization with *P. berghei* CS^5M^ parasites and analysis at effector (day 7) and memory (day 28) timepoints (Figure 7C). This protocol was designed to enable us to control for any unrelated differences between Ly5A and Ly5B mice.

**Figure 7.**
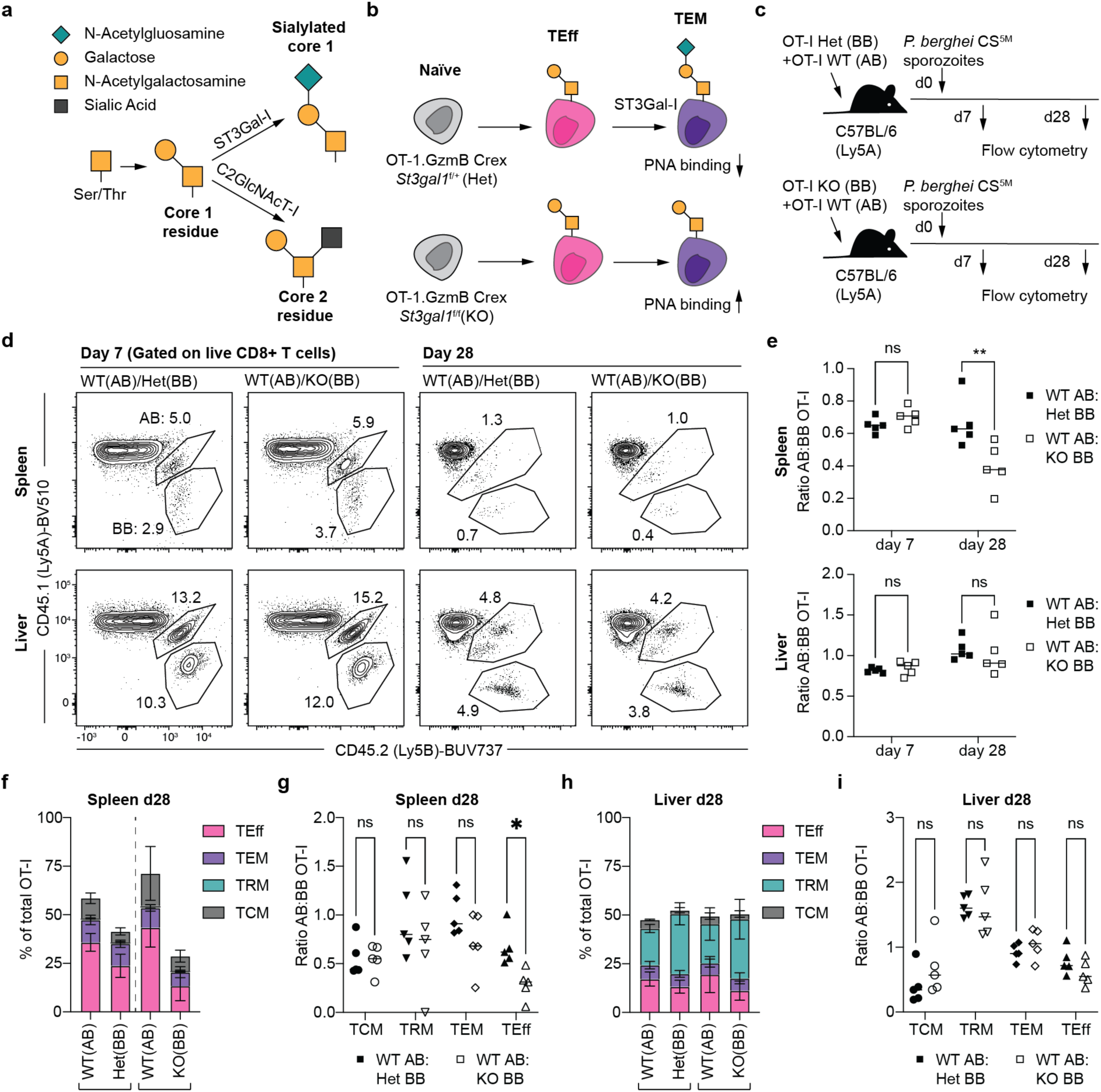
Forced expression of ASGPs reduces accumulation of effector CD8+ T cells in the spleen. (A) Graphical representation of the normal glycoprotein pathway of glycosylation modifications from a single Threonine/Serine residue to a Core-1 residue. Enzymatic modification via St3GalI adds a sialic acid residue to the terminal core 1 residue, whereas C2GlcNAcT-I adds a N-acetylglucosamine residue resulting in a core-2 residue with each residue having its own distinct, functional properties. (B) Rationale behind the generation and desired conditional knockout model of CD8+ effector cell loss of *ST3GalI* enzyme ultimately resulting in effector cells with desialylated residues on core-1 molecules, resulting in greater PNA binding. (C) WT mice received equal ratios of both activated WT OT-I CD8+ lymphocytes (Ly5AB) and either *St3GalI* heterozygous conditional knockouts (GzmB Crex *St3GalI*^*+/-*^), or homozygous mutant conditional knockouts (GzmB Crex *St3GalI*^*-/-*^). All mice were then immunised as per previous experiments using 5×10^3^ RAS and transferred cells quantified at both day 7 and day 28 post immunisation from the spleen and liver. (D) Flow cytometry analysis of WT OT-I cells (AB) and both homozygous and heterozygous *St3GalI* KO cells (BB) in the spleen and liver at day 7, and day 28 post immunisation. (E) Ratio of WT (AB) to GzmB Cre x *St3GalI* (BB) transgenic cells at day 7 and day 28 post immunisation in the spleen and liver. TEff, TEM, TRM and TCM cell phenotypes as a percentage of total OT-I cells and their WT to transgenic ratios recovered in the spleen (F-G) and liver (H-I) at day 28 post immunisation. Data are from a single experiment with 5 mice per group analyzed via 2-way ANOVA with Tukey post-test; bars are mean ± S.D; analyzed via LMM; * p<0.05, **p <0.01.

At day 7, both the Het and KO cells had a small survival disadvantage compared to control cells in both the spleen and liver, however the difference was not significant between these groups (Figure 6D-E). By day 28 however, the KO cells with enforced expression of ASGPs were lost from the spleen but not the liver (Figure 7D-E), consistent with our data using *in vitro* effector T cells, showing that ASGP expression is critical for interactions with the spleen but not the liver. Notably this overall loss of KO OT-I cells was driven by a loss of Teff cells in the spleen, but not other populations which was apparent in the spleen (Figure 7F-G) but not the liver (Figure 7H-I). Collectively these data suggest that high expression of ASGPs on Teff cells leads to accumulation in the liver, but also ultimate removal from the circulation by the spleen. In contrast TRMs are likely to downregulate these receptors to facilitate their maintenance in the liver.

## Discussion

Unlike lymphocytes in high endothelial venules, effector CD8+ T cell homing to the liver does not undergo selectin mediated rolling [46, 47], rather these cells crawl through the sinusoids, searching for antigen using an LFA-1-dependent patrolling behaviour [6, 7, 16]. It has been proposed in a model of HBV infection that, instead of rolling, CD8^+^ T cells in the liver initially tether to the endothelium via platelets [16]. We investigated this process in the context of *Plasmodium* infection and immunization. Using both thrombocytopenic (*Mpl*^*-/*^) mice and platelet-depleted recipient animals, we found that the large reduction in platelet concentration in circulation had at best minor effect on the effector function or accumulation of CD8 T-cells in the liver during *Plasmodium* infection. Platelet deficiency also did not affect the generation of liver TRMs in mice immunized with *Plasmodium* sporozoites.

Finally, we found that ASGP expression on the surface of activated CD8+ T cells did not mediate the accumulation of CD8+ T cells in the liver. Rather, ASGPs mediate effector CD8 T-cell accumulation in the red pulp of the spleen, where we hypothesize these cells are removed from the lymphocyte pool. Our results, thus, differ from the findings that platelets play important roles in CD8 T-cell homing in HBV model [16, 21].

Platelets have also been observed to play an important role in the accumulation of neutrophils cells in the liver in conditions of sterile injury [48]. We were able to observe some deficiency in CD8+ T cell-mediated killing in *Mpl*^-/-^ mice, however, this was unlikely to be due to thrombocytopenia as we were unable to see a similar effect in platelet-depleted animals. One limitation is that we do not know if our antibody depletion methods remove platelets that are already bound to activated CD8+ T cells. LFA-1 deficient (*Itgal*^-/-^) cells also showed a modest reduction of lymphocyte homing to the liver in platelet-deficient recipients. Collectively, these data suggest that LFA-1 is the dominant adhesion molecule involved in the retention of CD8+ T cells in the liver in our immunization models. One critical difference between our models and those used previously is the burden of antigen and inflammation in the liver: viral infection models induce a high burden of inflammation within the endothelial cells and parenchyma of the liver [16, 21], whereas the density of parasites in the *Plasmodium*-infected liver is low [49]. Roles for platelets in neutrophil accumulation in the liver have also been observed exclusively in conditions of inflammation [20, 48, 50].

One limitation of our studies is that all models of platelet deficiency have possible artifactual effects. *Mpl*^-/-^ mice have defects in haematopoiesis and retain around 15% of normal platelet numbers which may be sufficient for many functions [34, 39]. For example *Mpl*^-/-^ do not suffer from obvious bleeding problems [34]. Platelet depletion studies with antibodies can demonstrate systemic dysregulation of inflammatory processes, including changes to lymphocyte homing to the liver and spleen; notably, the use of anti-GPIbα antibodies results in the release of neuraminidase which may alter the sialylation of circulating and cell-associated proteins [44]. We hypothesise that this may in turn alter the homing of lymphocytes via the blockade of ASGPRs. This hypothesis would explain the apparent inhibition of effector CD8+ T cell homing to the spleen in antibody-mediated platelet-depletion models but not in *Mpl*^-/-^ mice.

ASGP exposure of the cell surface makes CD8+ T-cells vulnerable to apoptosis in the absence of antigen [15, 35]. The effects of desialylation on lymphocyte homing has been demonstrated with neuraminidase treated naïve cells, however these studies failed to investigate the role of ASGP expression during lymphocyte activation [13, 14]. The resultant enhanced binding to the liver was thought to be mediated by interactions with ASGPs and the ASGPR also known as the Ashwell-Morrell receptor [14, 51]. The ASGPR is abundantly expressed in the liver so the finding that ASGPs mediate accumulation in the spleen was surprising. It may be that cells accumulate in the spleen via interaction with a different receptor, one candidate would be Clec10a, which is abundantly expressed on macrophages and has been implicated in the clearance of desialylated platelets by Kupffer cells [52]. Importantly, our data suggest that the down regulation of ASGP expression on the cell surface may be required for the persistence of TCM and TRM in the spleen and liver, respectively.

Our study further supports the finding that LFA-1^hi^ CD8+ T cells in the liver represent a functional population capable of protecting against infection [6]. The formation of these protective populations in conditions of little or no inflammation does not require large numbers of platelets. Our data also suggests that it is the red pulp of the spleen, not the liver that is the true graveyard of senescent effector T cells. These data support an emerging paradigm that CD8+ T cells in the liver are a plastic population that not only protect against liver infection but also trans-differentiate into TRM populations in other tissues in the event of infection in other sites of the body [53].

## Acknowledgements

We thank M. Devoy, H. Vohra, and C. Gillespie of the Imaging and Cytometry Facility at the John Curtin School of Medical Research for assistance with flow cytometry and multiphoton microscopy.

## Supplementary Movie Captions

**Movie S1: Migration of naïve and activated CD8+ T cells in the liver** 2 x10^6^ SIINFEKL pulsed OT-I T-cells (CTV) and 2×10^6^ naïve GFP^+^ OT-I cells were transferred to C57BL/6 mice. 4 hours post transfer, livers of recipient mice were imaged using 2-photon microscopy with resonance scanning to collect time-lapse movement at 3 frames per second demonstrating elongated linear tracks of naïve cells (white) compared to short repetitive tracks of activated patrolling cells (blue). Scale bar = 50μm. Autofluorescence of liver stroma (green).

**Movie S2: Migration of CD8+ T cells in livers from wild-type and platelet-deficient mice** 2 x10^6^ SIINFEKL pulsed uGFP^+^ OT-I T-cells (green) were adoptively transferred to C57BL/6 and *Mpl*^*-/-*^ recipients. 4 hours post transfer, livers of recipient mice were prepared and imaged using 2-photon microscopy. For each series, a 50μm Z-stack (2μm/slice) was acquired using the galvo-scanner at a frame rate of ∼2 frames per minute demonstrating a liver sinusoidal crawling phenotype. Scale bar = 50μm. Autofluorescence of liver stroma (green).

