## Supplementary material for "Platelets are dispensable for the ability of CD8+ T cells to accumulate, patrol, kill and reside in the liver": Figure S1

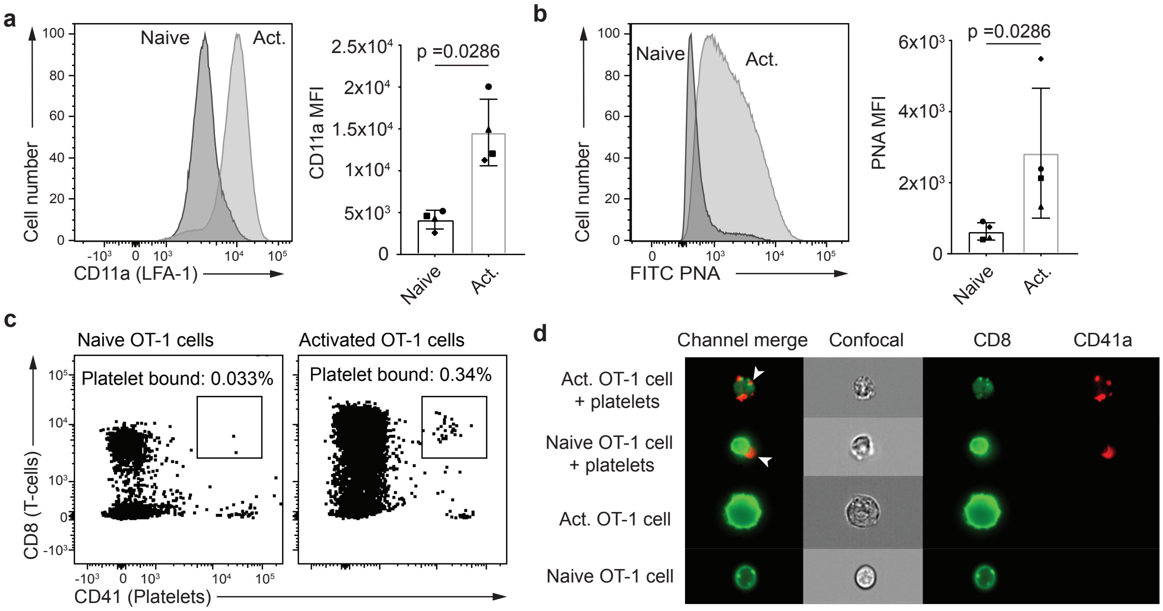


**Figure S1 *In vitro* activated CD8+ T-lymphocytes upregulate LFA-1, Asialylated terminal glycoprotein residues and platelet binding motifs.** (A) OT-I cells were activated *in vitro* and their expression of CD11a (LFA-1) was measured using flow cytometry. A representative histogram of CD11a expression on activated OT-I cells versus naïve OT-I cells (left) and a bar graph demonstrating CD11a mean fluorescence intensity (MFI) of 4 different samples pre- and post-activation; bars are mean ± S.D; analyzed by Mann-Whitney U test. (B) Activated OT-I PNA binding MFI from a representative mouse compared to a naïve cell sample (left) and a bar graph demonstrating PNA MFI of 4 different samples pre- and post-activation; bars are mean ± S.D; analyzed by Mann-Whitney U test. (C) Naïve and activated CD8+ lymphocytes were co-cultured with purified murine platelets and then stained for anti-CD8α and anti-CD41 using flow cytometry. Double positive cells indicated platelet bound CD8+ T-cells. (D) Cells stained for flow cytometry were analysed using IVIS spectrum flow cytometry to visualise binding interactions (arrow) between naïve and activated cells (CD8-green) to platelets (CD41-red).
