## Supplementary material for "Platelets are dispensable for the ability of CD8+ T cells to accumulate, patrol, kill and reside in the liver": Figure S2

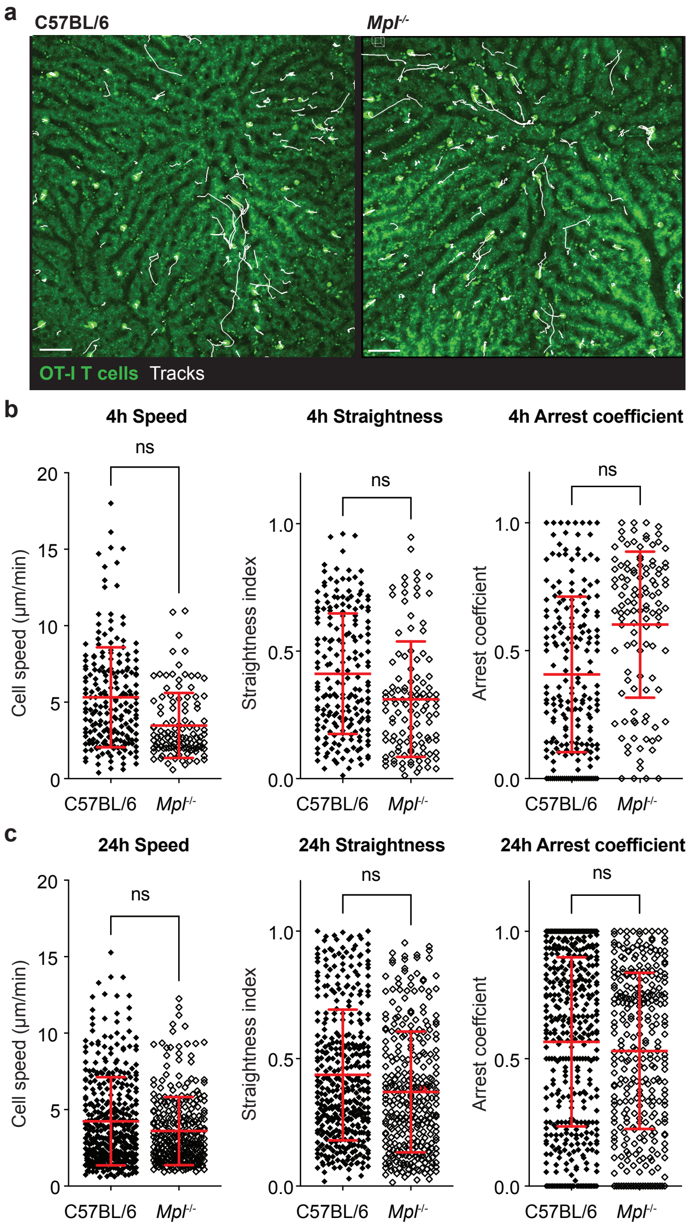


**Figure S2 *In vivo* activated OT-I cells demonstrate delayed crawling behaviour in the liver in *Mpl*$^{-/-}$ recipients compared to WT.** (A) uGFP-OT-I cells were activated *in vitro* and adoptively transferred into either C57BL/6 mice or platelet deficient *Mpl*$^{-/-}$ recipients. At 4 hours post transfer, the livers underwent intravital imaging using a Galvano scanner at a depth of 50μm and a frequency of ~2 frames per minute, the tracks of each cell are represented by white lines. (B) Imaging was analysed using IMARIS software to track each cell and measure the mean speed, track straightness and arrest coefficient (time spent arrested) of cells at 4 hours post transfer and 24 hours post transfer. Scale bar = 50μm. Data are pooled from 3-4 mice per condition analyzed via LMM.
