## Supplementary material for "Platelets are dispensable for the ability of CD8+ T cells to accumulate, patrol, kill and reside in the liver": Figure S3

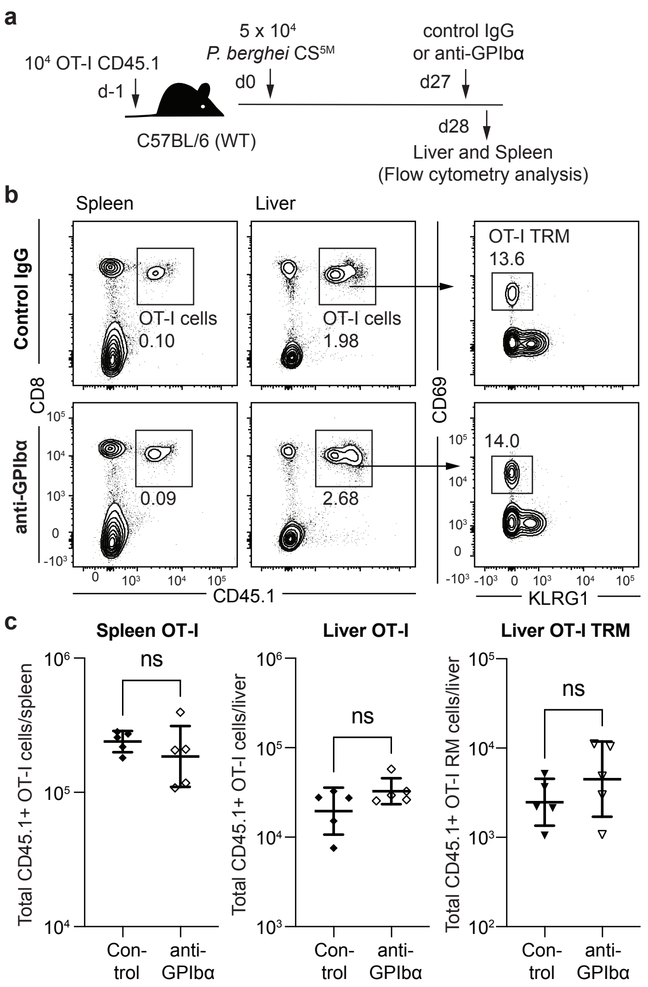


**Figure S3 *In vivo* generation of tissue resident memory T cells is not affected by the absence of platelets in the spleen and liver.** (A) C57BL/6 recipients received 5x10$^4$ OT-I cells prior to immunisation with 5x10$^4$ *P.berghei* CS$^{5M}$ RAS 24h later to generate CD8 tissue resident memory (TRM) populations *in vivo* after 28 days. 1 day prior to analysis (d27) mice were treated with anti-GPIbα to deplete circulating platelets. Total transferred cells recovered from the spleen and liver were assessed at day 28 post immunisation from both C57BL/6 and anti-GPIbα-treated recipients. (B) Representative flow cytometry plots and (C) summary data pooled from 2 independent experiments each with 5 mice per group and analyzed via LMM; bars are mean ± S.D.
