## Supplementary material for "Platelets are dispensable for the ability of CD8+ T cells to accumulate, patrol, kill and reside in the liver": Figure S4

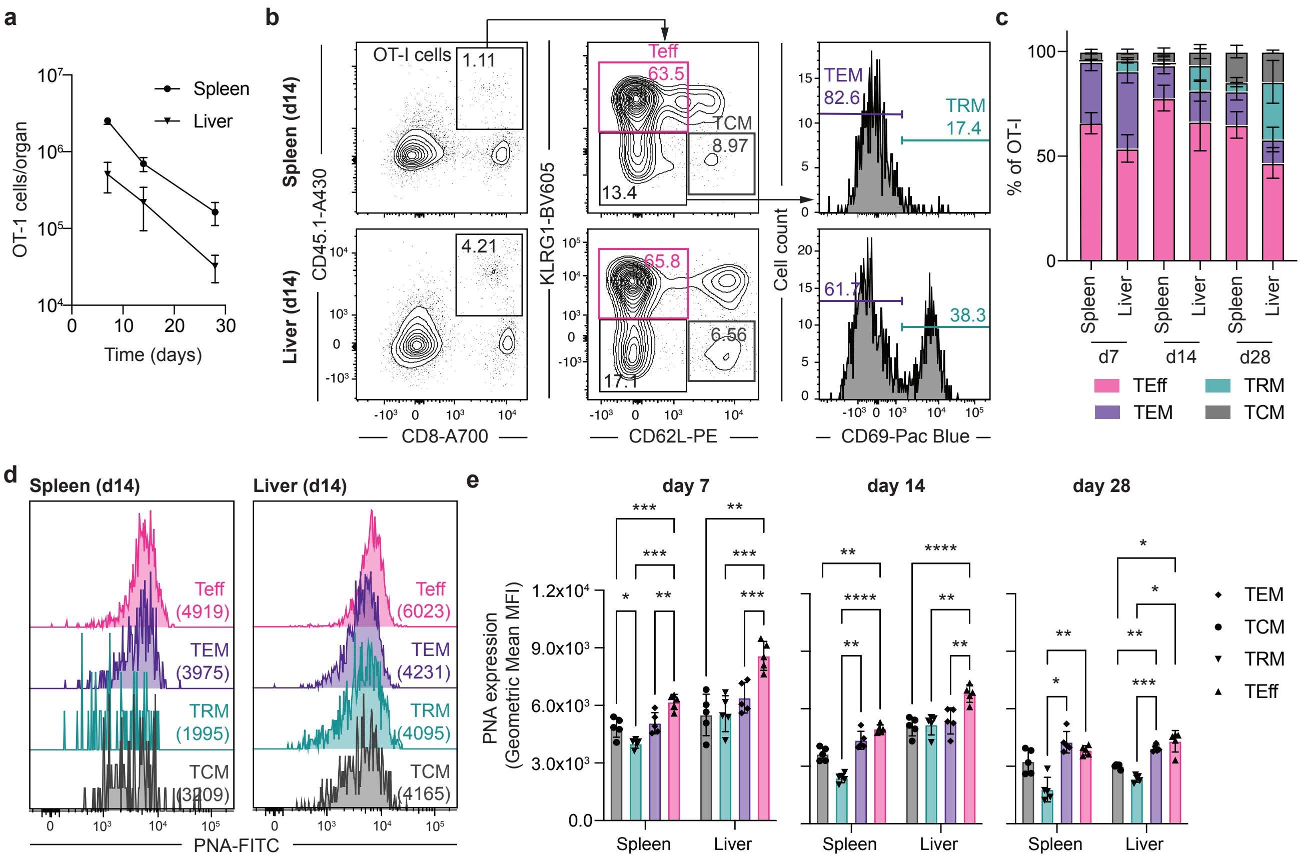

**Figure S4 Lectin binding motifs on lymphocyte glycoproteins are upregulated in the liver and spleen during effector cell differentiation *in vivo*.** (A) C57BL/6 recipients received 5x10$^4$ OT-I cells prior to immunisation with 5x10$^4$ *P.berghei* CS$^{5M}$ RAS 24h later to generate CD8+ effector (TEff), effector memory (TEM), central memory (TCM) and Tissue resident memory (TRM) populations *in vivo* over 28 days. Total transferred cells recovered from the spleen and liver were assessed at day 7, day 14 and day 28 post immunisation. (B) Cells were then taken from the spleen and liver at each timepoint and quantified from total recovered transgenic cells using flow cytometry; TEff - KLRG1$^{hi}$CD62L$^{lo}$ (pink), TCM – KLRG1$^{lo}$CD62L$^{hi}$ (grey), TEM -KLRG1$^{lo}$CD62L$^{lo}$CD69$^{-}$ (purple) and TRM - KLRG1$^{lo}$CD62L$^{lo}$CD69$^{+}$ (green). (C) Transgenic cell phenotypes were quantified as a percentage of recovered OT-I cells at day 7, day 14 and day 28. (D) The expression of desialylated glycoprotein binding motifs in each cell phenotype was quantified using the FITC conjugated lectin, Peanut agglutinin (PNA). (E) Mean fluorescence intensity of FITC conjugated PNA in each transgenic CD8 phenotype at day 7, day 14 and day 28 post immunisation. Results are from a single experiment; bars are mean ± S.D; analyzed via 2-way ANOVA with Tukey's post-test; * p<0.05, **P<0.01, ***p<0.001, ****p<0.0001.
